# HOIL-1-catalysed ubiquitylation of unbranched glucosaccharides and its activation by ubiquitin oligomers

**DOI:** 10.1101/2021.09.10.459791

**Authors:** Ian R. Kelsall, Elisha H. McCrory, Yingqi Xu, Cheryl L. Scudamore, Sambit K. Nanda, Paula Mancebo-Gamella, Nicola T. Wood, Axel Knebel, Stephen J. Matthews, Philip Cohen

## Abstract

HOIL-1, a component of the Linear Ubiquitin Assembly Complex (LUBAC), ubiquitylates serine and threonine residues in proteins, forming ester bonds (Kelsall et al, 2019, PNAS 116, 13293-13298). Here we report that mice expressing the E3 ligase-inactive HOIL-1[C458S] mutant accumulate polyglucosan in brain, cardiac muscle and other organs, indicating that HOIL-1’s E3 ligase activity is essential to prevent these toxic polysaccharide deposits from accumulating. We found that HOIL-1 monoubiquitylates glycogen and α1:4-linked maltoheptaose *in vitro* and identify the C6 hydroxyl moiety of glucose as the site of ester-linked ubiquitylation. The HOIL-1-catalysed monoubiquitylation of maltoheptaose was accelerated >100-fold by Met1-linked or Lys63-linked ubiquitin oligomers, which interact with the catalytic RBR domain of HOIL-1. HOIL-1 also transferred preformed ubiquitin oligomers to maltoheptaose *en bloc*, producing polyubiquitylated maltoheptaose in one catalytic step. The Sharpin and HOIP components of LUBAC, but not HOIL-1, bound to amylose resin *in vitro*, suggesting a potential function in targeting HOIL-1 to unbranched glucosaccharides in cells. We suggest that monoubiquitylation of unbranched glucosaccharides may initiate their removal by glycophagy to prevent precipitation as polyglucosan.

## Introduction

Mutations that reduce the expression of HOIL-1 (haem-oxidised IRP2 ubiquitin ligase-1), also called RBCK1 (RING-B-Box-coiled-coil protein interacting with PKC 1 or RANBP2-Type and C3HC4-type zinc finger-containing protein 1) cause both auto-inflammation and immuno-insufficiency in humans (Boisson *et al*, 2012; Phadke *et al*, 2020) and immuno-insufficiency in mice (MacDuff *et al*, 2015; Tokunaga *et al*, 2009). However, HOIL-1 deficiency in humans also leads to cardiomyopathy and death from heart failure in early adulthood (Boisson *et al*., 2012; Fanin *et al*, 2015; Krenn *et al*, 2018; Nilsson *et al*, 2013; Phadke *et al*., 2020; Wang *et al*, 2013), which is unrelated to the immune defects, and arises from the progressive accumulation of toxic polyglucosan bodies in cardiac muscle and other tissues, such as the brain, with some patients also displaying cognitive impairment (Chen *et al*, 2021; Phadke *et al*., 2020). Mice expressing low levels of HOIL-1 (Fujita *et al*, 2018) also form toxic polyglucosan bodies in cardiac muscle (MacDuff *et al*., 2015), brain and spinal cord (Sullivan *et al*, 2018), but it is the brain that is affected predominantly in mice, the animals displaying defects in learning, memory and motor coordination (Sullivan *et al*., 2018).

Polyglucosan bodies are dense inclusions of starch-like polysaccharide that are insoluble because they lack the α1:6 branch points found in the glucose polymer glycogen. Consequently, they have defective metabolism compared to glycogen and are resistant to digestion with α-amylases (Hedberg-Oldfors & Oldfors, 2015). The frequency of the α1:6 branch points, which occur after about every twelve α1:4-linked glucose units, determines the topology, structure, and solubility of glycogen. Mutations in glycogen metabolising enzymes cause a variety of glycogen storage diseases characterised by aberrant glycogen deposits (Kanungo *et al*, 2018).

HOIL-1, is a component of the linear ubiquitin chain assembly complex, LUBAC (Dittmar & Winklhofer, 2019; Kirisako *et al*, 2006; Liu & Pan, 2018). It is well established that this trimeric complex, comprising HOIL-1, HOIP (HOIL-1 interacting protein) and Sharpin (Shank-associated RH domain interactor) is required in signal transduction pathways that generate inflammatory mediators or lead to cell death (Gerlach *et al*, 2011; Ikeda, 2015; Ikeda *et al*, 2011; Sasaki & Iwai, 2015; Shimizu *et al*, 2015; Tokunaga *et al*, 2011). In these pathways, Met1-linked ubiquitin (M1-Ub) oligomers generated by the E3 ligase HOIP interact with several proteins that regulate innate immunity, such as the NEMO regulatory subunit of the canonical IκB kinase (IKK) complex. This leads to IKK activation and the phosphorylation and activation of its substrates, which include the transcription factors NF-κB and IRF5 (Lopez-Pelaez *et al*, 2014; Ren *et al*, 2014; Scheidereit, 2006) that stimulate expression of the mRNAs encoding many inflammatory mediators.

Like HOIP, HOIL-1 is a member of the “RING in-between RING” (RBR) subfamily of E3 ubiquitin ligases but, unlike HOIP, it does not generate Met1-linked ubiquitin oligomers. Instead, it catalyses the attachment of ubiquitin to serine and threonine residues in proteins, forming ester bonds, and its substrates include the IRAK1 and IRAK2 components of Myddosomes that have critical roles in inflammatory mediator production (Kelsall *et al*, 2019). The failure to generate ester-linked ubiquitin chains in knock-in mice expressing the E3 ligase-inactive HOIL-1[C458S] mutant can enhance or reduce the production of inflammatory cytokines, depending on the ligand, receptor and immune cell type (Petrova *et al*, 2021).

Since HOIL-1 ubiquitylates the hydroxyl side chains of serine and threonine residues (Kelsall *et al*., 2019; Rodriguez Carvajal *et al*, 2021), we hypothesized that it might also ubiquitylate the hydroxyl groups in glucose and so be able to ubiquitylate glycogen directly. Here, we report that HOIL-1 does indeed ubiquitylate glycogen and the smaller model oligosaccharide maltoheptaose *in vitro* and identify the site of ubiquitylation. Based on these and other unexpected findings reported in this paper, we propose a new role for HOIL-1 and LUBAC in the ubiquitylation of unbranched glycogen molecules that may initiate their elimination from cells before they precipitate as toxic polyglucosan deposits.

## Results

### Polyglucosan accumulates in the brain and heart of HOIL-1[C458S] mice

HOIL-1 knockout (KO) mice exhibit early embryonic lethality (Fujita *et al*., 2018; Peltzer *et al*, 2018), but HOIL-1 “KO” mice expressing low levels of a truncated protein comprising the N-terminal Ubiquitin-Like (UBL) and Npl4 zinc finger (NZF) domains, but lacking the C-terminal region including the catalytic RBR domain (Fujita *et al*., 2018), accumulate polyglucosan in their brain, spinal cord and heart (MacDuff *et al*., 2015; Sullivan *et al*., 2018). Because HOIL-1 stabilises its binding partner HOIP, the expression of HOIP and Sharpin are also greatly reduced in these HOIL-1 “KO” mice. It was therefore unclear whether the accumulation of polyglucosan was caused by the loss of the E3 ligase activity of HOIL-1, the loss of the E3 ligase activity of HOIP or reduced expression of the non-catalytic domains of HOIL-1, HOIP or Sharpin. In contrast, HOIL-1, HOIP and Sharpin are expressed at normal levels in the E3 ligase-inactive HOIL-1[C458S] knock-in mice that we have described previously (Kelsall *et al*., 2019). We therefore investigated whether polyglucosan was present in the tissues of these mice. We found that polyglucosan did indeed accumulate in the brain, heart and other tissues of the HOIL-1[C458S] mice, but not in their wild-type littermates. Particularly high levels of polyglucosan were present in the hind brain (pons) and the hippocampus, the extent of deposition being similar in mice varying in age from 0.5-1.5 years (**Fig. 1A-D**). Polyglucosan was present in lower amounts in the brain cortex (**Figs. EV1A and EV1B**). It was also present in small amounts in the heart, reaching a maximum level after about a year (**Fig. 1E and 1F**), as well as in the lungs and liver (**Figs. EV1C-F**). These results establish that the E3 ligase activity of HOIL-1 is required to prevent the accumulation of polyglucosan in several tissues.

**Figure 1.**
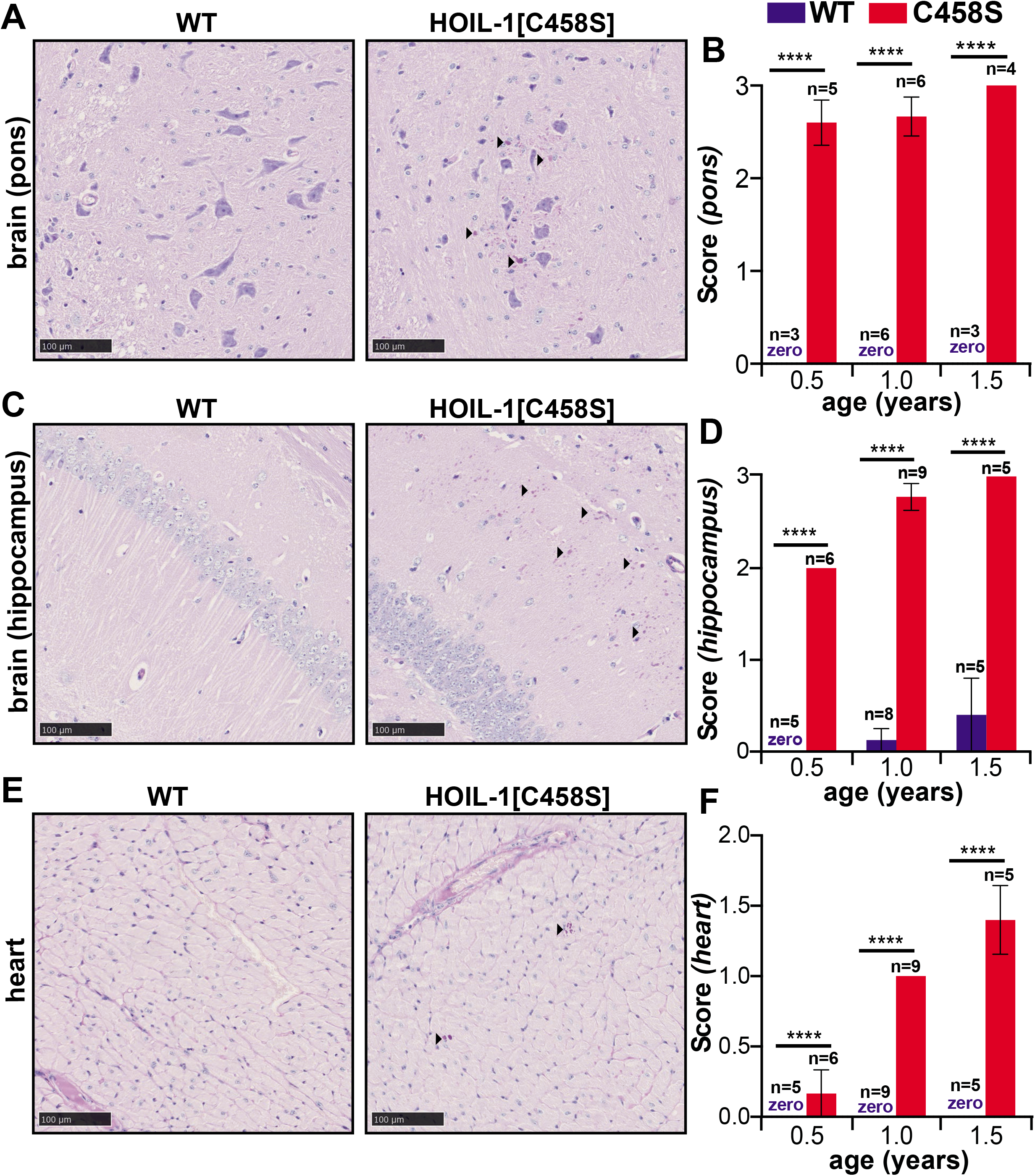
Deposition of α-amylase-resistant polyglucosan deposits in the hind brain and heart of HOIL-1[C458S] mice. **(A, C, E)** Representative PAS-stained sections of the pons (A) and hippocampal (C) regions of the brain and heart (E) of one year old HOIL-1[C458S] and WT mice are shown. Scale bar = 100 μm. Arrow heads indicate α-amylase-resistant PAS-positive polyglucosan deposits. **(B, D, F)** Graphs showing α-amylase resistant PAS scores of the pons (B) and hippocampal (D) regions of the brain and the heart (F) of HOIL-1[C458S] (red) and WT (blue) mice aged 0.5, 1.0 and 1.5 years. The number of biological replicates analysed at each age is indicated. The word zero highlighted in blue indicates that no α-amylase resistant, PAS-positive material could be detected in the WT mice. The error bars show mean ± SEM. Statistical significance between the genotypes was calculated by using two-way ANOVA and Šidák’s multiple comparison’s test. **** denotes p<0.0001.

### HOIL-1 ubiquitylates glycogen *in vitro*

We considered how HOIL-1 might prevent the deposition of polyglucosan and wondered whether HOIL-1 might not only ubiquitylate the hydroxyl side chains of serine and threonine residues in proteins, but also the hydroxyl moieties present in glucose. To investigate whether HOIL-1 was capable of ubiquitylating glycogen directly we initially used an *in vitro* fluorescence-based assay in which commercially available bovine liver glycogen was incubated with HOIL-1, E1, E2, Mg^2+^-ATP and fluorescent Cy5-labelled ubiquitin. Following SDS-PAGE, the glycogen (detected by Periodic acid-Schiff [PAS] staining), is too large to enter the gel and remains at the origin **(Fig. 2A)**, as does a portion of the fluorescent Cy5-labelled ubiquitin **(Fig. 2B, lanes 5 and 6)**. The comigration of Cy5-labelled ubiquitin and glycogen required the presence of every reaction component. This high molecular mass Cy5-ubiquitin signal was lost if the reaction was incubated with α-amylase to degrade the glycogen **(Fig. 2B, lanes 7 and 8)** or incubated with the oxyester-specific nucleophile hydroxylamine **(Fig. 2B, lanes 9 and 10)**. These experiments indicated that HOIL-1 had ubiquitylated glycogen and that ubiquitin was attached to it via an oxyester bond.

**Figure 2.**
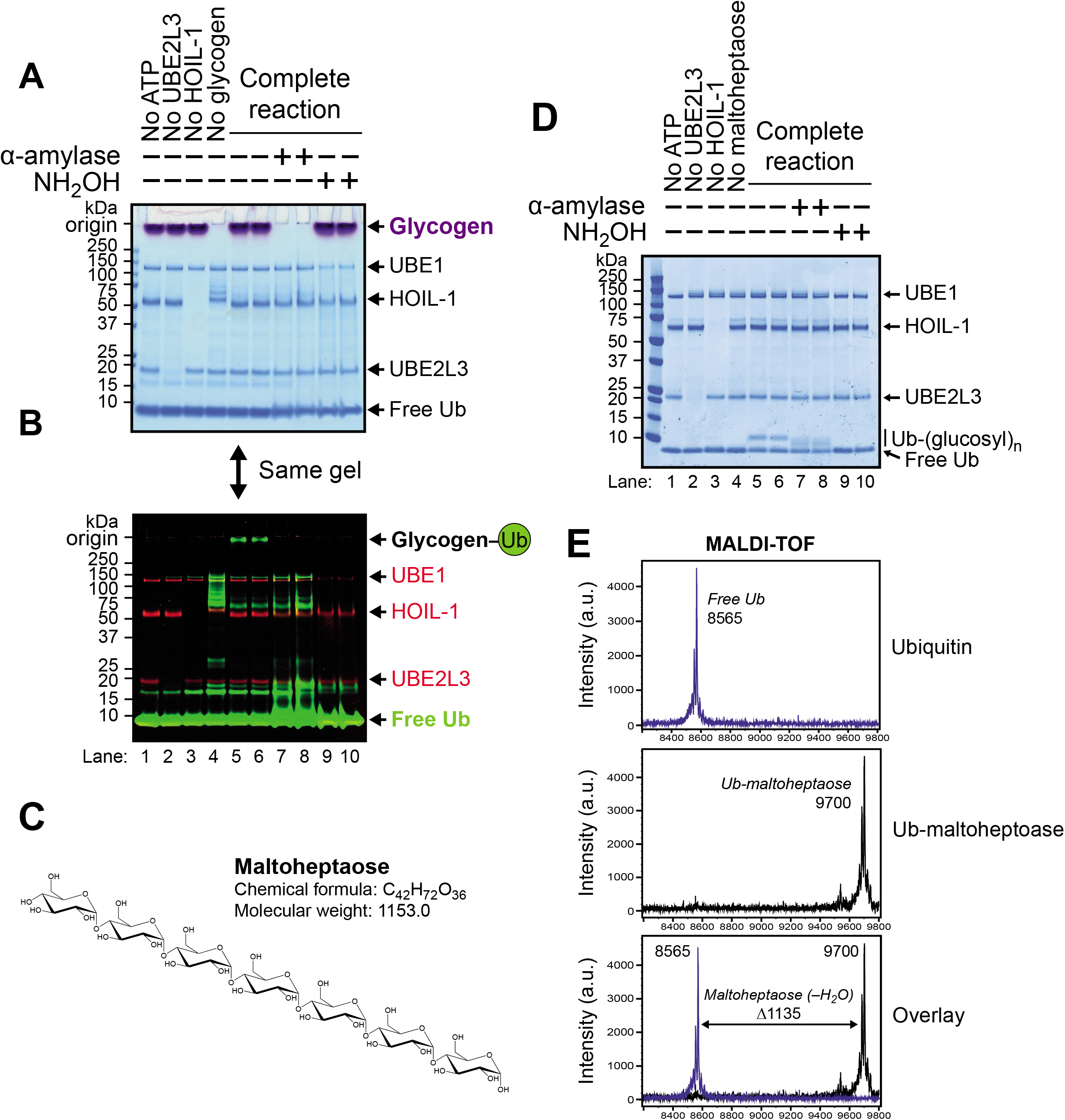
HOIL-1 ubiquitylates oligo- and polysaccharides *in vitro*. **(A)** HOIL-1 activity towards purified bovine liver glycogen was measured using Cy5-labelled ubiquitin. Control assays lacking ATP, UBE2L3, HOIL-1 and glycogen are indicated, as are subsequent treatments with 50 μg/mL human salivary α-amylase and 1.5 M hydroxylamine (NH_2_OH). The gel has been stained using the periodic acid-Schiff (PAS) method to visualise glycogen (purple-magenta) and Coomassie Blue stain to visualise protein. **(B)** Prior to PAS and Coomassie staining, the gel shown in (A) was stained with Flamingo fluorescent protein stain (shown in red) and glycogen ubiquitylation observed by visualising the fluorescent Cy5-ubiquitin signal (shown in green). **(C)** Chemical structure of the α-1,4-linked heptasaccharide maltoheptaose. **(D)** *In vitro* ubiquitylation of maltoheptaose by HOIL-1. Assays were treated with α-amylase or hydroxylamine as indicated. Reaction products were detected by Coomassie staining. **(E)** Ubiquitylated maltoheptaose was purified from reaction components and analysed by MALDI-TOF, revealing an additional 1135 Da attached to the ubiquitin. This is consistent with addition of maltoheptaose to ubiquitin via an oxyester linkage (expected additional mass is 1153 for maltoheptaose minus 18 to account for the loss of a water molecule during ester bond formation).

### Small malto-oligosaccharides are mono-ubiquitylated by HOIL-1 *in vitro*

The majority of the glucose units in glycogen are linked via α-1,4-glycosidic bonds. The linear oligosaccharide maltoheptaose contains seven glucose units linked by such bonds **(Fig. 2C)** and was used as a simple model substrate. The replacement of glycogen by maltoheptaose in the *in vitro* ubiquitylation reaction led to the formation of a single more slowly migrating form of ubiquitylated maltoheptaose in a concentration and time-dependent manner **(Figs. EV2A and EV2B)**. As observed for glycogen, this adduct was sensitive to treatment with both hydroxylamine and α-amylase, although the ubiquitin was still bound to maltose and maltotriose after digestion with α-amylase, because complete hydrolysis to glucose did not take place (Zakowski & Bruns, 1985) **(Fig. 2D, lanes 7 and 8)**. Mass spectrometry analysis of ubiquitylated-maltoheptaose established that it was a monoubiquitylated species (**Fig. 2E**).

### NMR Spectroscopy identifies the hydroxyl group attached to glycosyl carbon C6 as the site of attachment of ubiquitin

To identify the ubiquitin ligation site, a hybrid labelled sample was prepared in which ubiquitin was [^15^N,^13^C]-labelled and ligated to unlabelled maltoheptaose. The purpose was to reassign ubiquitin and measure scalar couplings or nuclear Overhauser effects and connectivities across the protein-carbohydrate interface. As anticipated, the ^1^H-^15^N HSQC spectrum of [^15^N,^13^C] ubiquitin-maltoheptaose shows the characteristic peak patterns of ubiquitin and any peak movements could be reassigned readily. Two major amide chemical shifts differences can be observed at the C-terminus, namely G75 and G76, when compared to free ubiquitin (**Fig. 3A**). Most notably, the amide ^15^N and ^1^HN signals of G76 are shifted upfield by −5.4 ppm and downfield by 0.3 ppm, respectively, which is consistent with the average C-terminal amide values when the charged carboxylate is removed by modification (Ulrich *et al*, 2008). Taken together with the observation that amide linewidths are comparable to free ubiquitin, these data indicate that a single ubiquitin moiety is conjugated to the oligosaccharide via its C-terminal glycine residue. Interestingly, the C-^15^N and ^1^HN signals of G76 exhibit multiple chemical shift environments, with at least four resolved positions, suggesting different isomers of the conjugate, with individual maltoheptaose molecules being singly ubiquitylated at distinct sites along the oligosaccharide.

**Figure 3.**
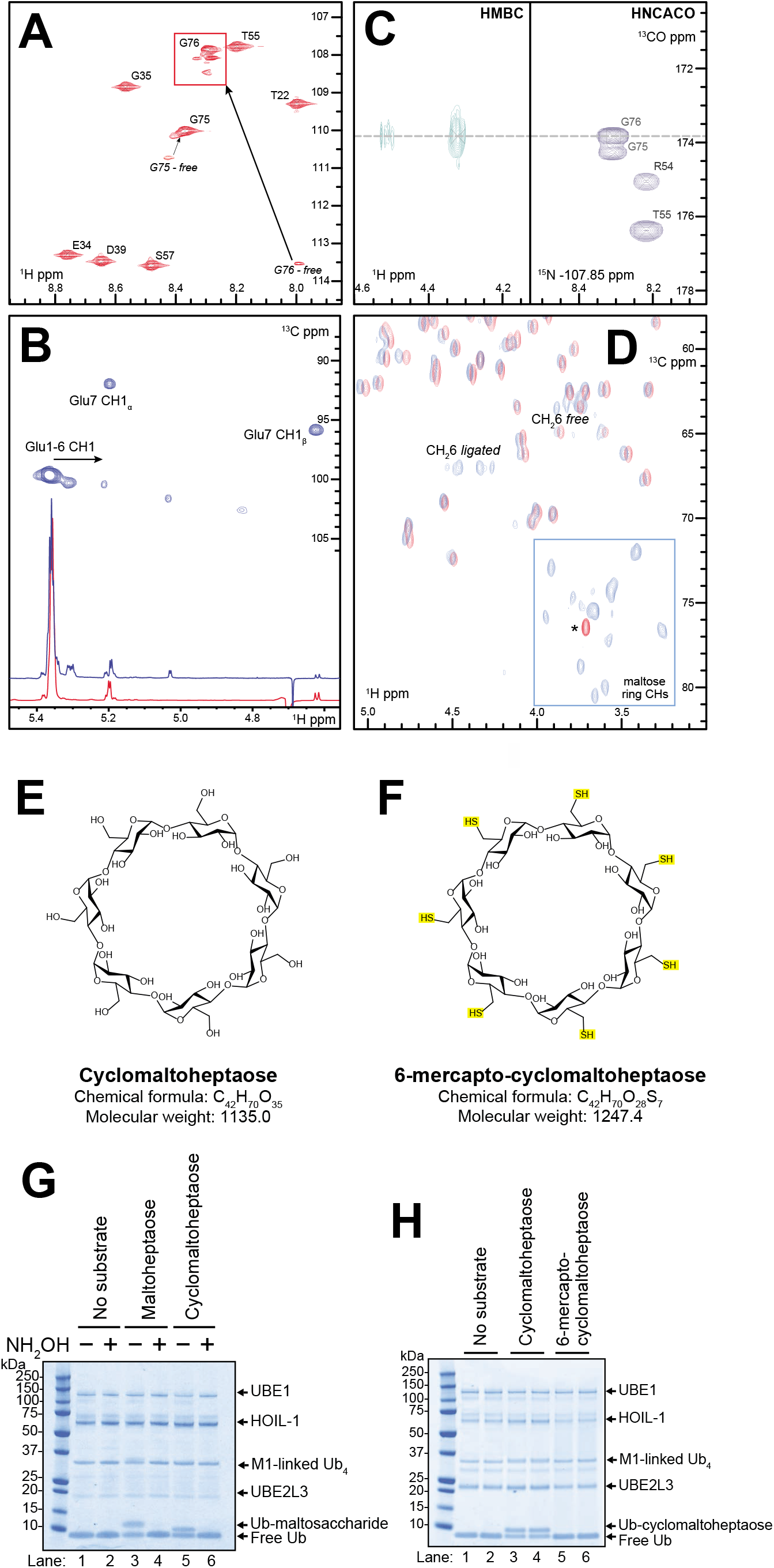
NMR spectroscopy identifies the hydroxyl groups ubiquitylated by HOIL-1 in maltoheptaose and maltose. **(A)** Region from the ^1^H-^15^N HSQC spectrum of [^15^N,^13^C] Ub-maltoheptaose with key protein amide assignments labelled. Arrows indicate chemical shift changes dues to C-terminal conjugation. (**B**) 1D ^1^H NMR overlay for the anomeric region of maltoheptaose (red) and [^15^N,^13^C] Ub-maltoheptaose (blue) together with the ^1^H-^13^C HSQC of the latter. Key anomeric assignments labelled, and arrow indicate chemical shift differences. (**C**) ^1^H-^13^C HMBC alongside HNCACO showing 3-bond connectivity from G76 carbonyl to the 6-CH2 chemical shifts. (**D**) Overlay of the ^1^H-^13^C HSQC spectra for free ubiquitin (red) and Ub-maltose (blue). Assignment for ligated and free 6-CH2 position of the disaccharide. Asterisk indicates a buffer component. **(E)** Chemical structure of cyclomaltoheptaose (β-cyclodextrin). **(F)** Chemical structure of 6-mercapto-cyclomaltoheptaose (heptakis-(6-deoxy-6-mercapto)-β-cyclodextrin). **(G)** HOIL-1 activity against the indicated maltosaccharides (2 mM) was assayed and visualised by staining with Coomassie protein stain. Where indicated assays were treated with 1.5 M hydroxylamine. **(H)** *In vitro* ubiquitylation of the indicated cyclodextrins (2 mM) by HOIL-1, visualised by Coomassie staining.

Comparison of the anomeric region of the 1D ^1^H NMR spectrum for the ubiquitylated maltoheptaose with free maltoheptaose (**Fig. 3B**), shows that while the anomeric protons of the α and β reducing end sugar are largely unaffected, there are a number of significant chemical shift changes for the anomeric protons of the other glucose residues. As the anomeric position of these sugar residues cannot be targeted by the E3 ligase, the chemical shift changes reflect the effects of ubiquitylation on other positions within these sugars.

To identify unambiguously which position(s) on the glucose units were modified with ubiquitin, we exploited the hybrid labelling scheme of the protein-carbohydrate conjugate and used an ^1^H-^13^C HMBC experiment to detect the 3-bond scalar coupling between the sugar protons and the ^13^C carbonyl of G76. These ^3^*J*_COCH_ couplings are typically 2-3 Hz and correlations in the HMBC would reveal the specific ligated positions. Only one significant correlation could be observed at 4.30 and 4.50 ppm (**Fig. 3C**), corresponding to two protons of a CH_2_ group, which could be readily assigned using ^1^H correlation NMR spectroscopy to the sidechain 6-CH_2_ position of the glucose residues.

To confirm these assignments, we prepared an unlabelled version of ubiquitylated maltose and compared the ^1^H-^13^C HSQC spectrum with that for free ubiquitin (**Fig. 3D**). The conjugated 6-CH_2_ group is clearly identifiable at 4.30 and 4.50 ^1^H ppm (67.0 ^13^C ppm) and can be assigned to the non-reducing glucose unit. These chemical shifts match those for 6-CH_2_ groups typically found in (α1-6) glucose chains (Dobruchowska *et al*, 2012), thereby confirming the 6-position modification. We also note a minor set of proximal CH_2_ peaks in this region of the spectrum, which could either represent a different conformation of the non-reducing 6-CH_2_ group due to restriction in free bond rotation from the attached ubiquitin or some ligation to the reducing end sugar 6-CH_2_ group, although there are no apparent chemical shift changes for the anomeric proton at the reducing end (**Fig. 3B**). Peaks for a second 6-CH_2_ group are also observed at 3.85 and 3.90 ^1^H ppm (63.5 ^13^C ppm) chemical shifts, which correspond to an unmodified glucose 6-CH_2_OH group (Bekiroglu *et al*, 2003) and this belongs to the reducing end carbohydrate residue of maltose. The ring CH protons are clearly visible between 71 and 81 ppm and these show no significant chemical shift changes when compared to free maltose, suggesting that any ubiquitylation of the ring CHOH groups is absent, or is minimal and beyond detection here.

Consistent with the NMR analysis we found that cyclomaltoheptaose lacking the reducing C1 hydroxyl and non-reducing C4 hydroxyl groups could still be ubiquitylated **(Figs. 3E and 3G)**, but the 6-mercaptan derivative of cyclomaltoheptaose lacking any 6-CH_2_OH groups could not **(Figs. 3F and 3H**).

It should be noted that the hydroxymethyl (CH_2_-OH) group of a glycosyl unit is identical to the hydroxymethyl side-chain of the amino acid serine, a previously identified target for HOIL-1 ubiquitylation (Kelsall *et al*., 2019).

### HOIP and Sharpin bind to amylose resin

Since the rate of ubiquitylation of glycogen was slow, taking many hours to convert most of the ubiquitin in the assay to monoubiquitylated maltoheptaose (**Fig. EV2B**), we wondered whether a mechanism(s) might be needed to accelerate the HOIL-1-catalysed ubiquitylation of glycogen in cells, if this reaction was to be of physiological significance. We therefore initially investigated the binding of HOIL-1 and other proteins to the α1:4-linked oligosaccharide amylose to evaluate their glucan-binding properties (**Fig. 4A**). Unlike maltose/maltodextrin binding protein (MBP), HOIL-1 did not bind to amylose-agarose resin, similar to other negative control proteins, such as glutathione S-transferase (GST). However, to our surprise, the other two components of LUBAC, Sharpin and HOIP, both bound to amylose resin, the latter behaving similarly to MBP in the pull-down assays (**Fig. 4A**). This suggested that one role for the interaction of HOIL-1 with HOIP and Sharpin in LUBAC may be to enable HOIP and/or Sharpin to target HOIL-1 to glycogen in cells.

**Figure 4.**
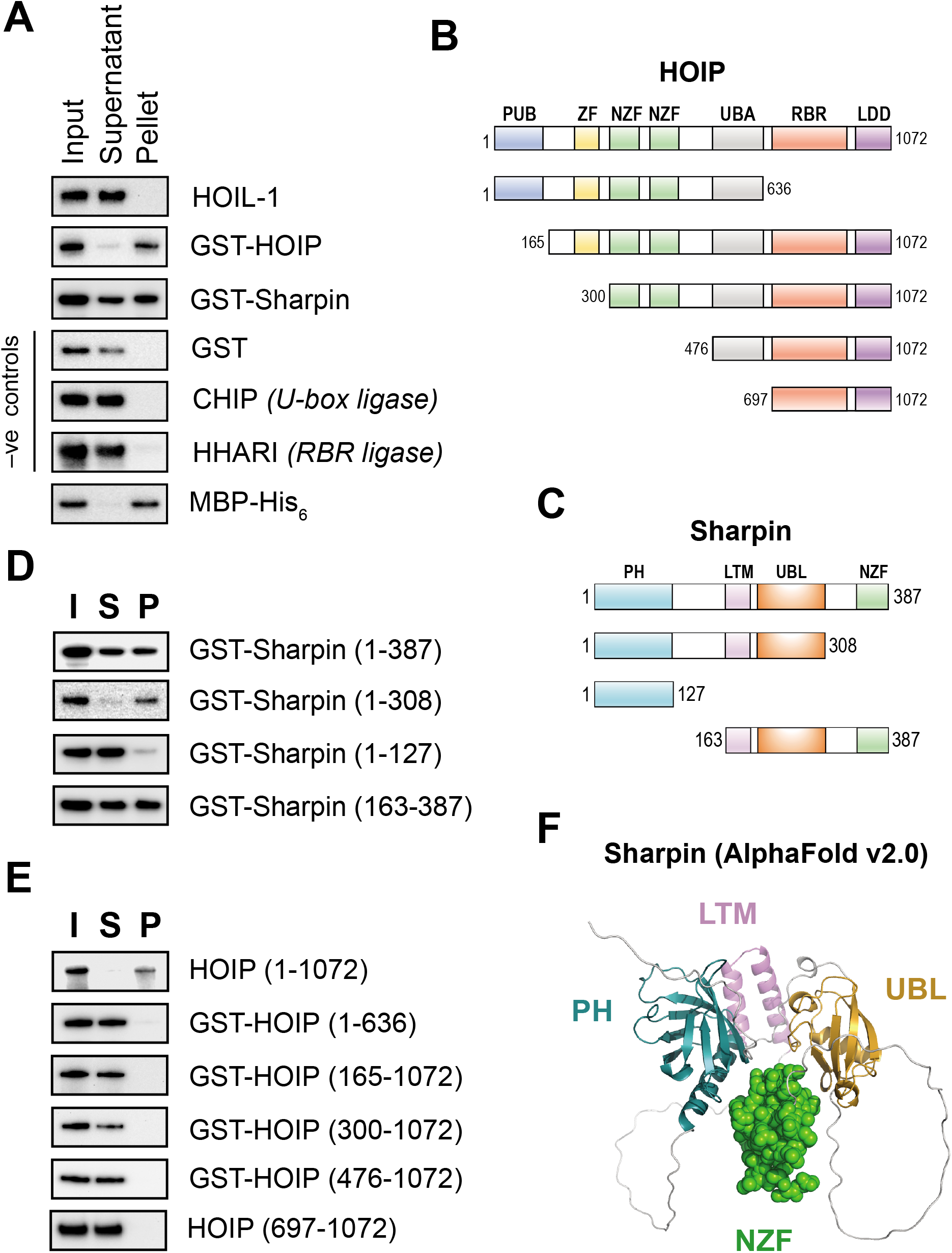
HOIP and Sharpin bind amylose *in vitro*. **(A)** 1 μg of each of the indicated recombinant proteins was incubated with amylose resin for 60 min, the beads were pelleted by centrifugation, and proteins in the pellet (P) and supernatant (S) were compared to the protein input (I) by means of western blotting with the appropriate antibodies. **(B)** Schematic representation of the truncated HOIP constructs used in (E). Images not drawn to scale. PUB, peptide N-glycanase and UBA- and UBX-containing protein domain; ZF, zinc finger domain; NZF, Npl4 zinc finger domain; UBA, ubiquitin-associated domain; RBR, RING-in-between-RING domain; LDD, linear ubiquitin chain determining domain. **(C)** Schematic of Sharpin constructs used in (D). Images not shown to scale. PH, pleckstrin homology domain; LTM, LUBAC-tethering motif; UBL, ubiquitin-like domain. **(D)** Amylose binding assays of full-length (1-387) and truncated Sharpin mutants, visualised by immunoblotting. **(E)** Amylose binding assays of full-length (1-1072) and truncated HOIP protein. **(F)** The structure of human Sharpin was predicted using AlphaFold v2.0. The PH, LTM and UBL domains are shown as ribbons, the NZF domain is depicted as green spheres.

There are no obvious carbohydrate-binding domains in the primary sequence of either HOIP or Sharpin (**Figs. 4B and 4C**). We therefore analysed various protein truncations for their ability to bind amylose in our pull-down assay (**Figs. 4D and 4E)**. The isolated N-terminal Pleckstrin Homology (PH) domain of Sharpin did not bind to amylose resin, but removal of Sharpin’s Npl4 zinc finger (NZF) domain actually increased amylose binding (**Fig. 4D**). None of the truncated HOIP constructs bound to amylose, implying that noncontiguous regions most likely contribute to glucan binding (**Fig. 4E**).

Recently, DeepMind and EMBL’s European Bioinformatics Institute created the AlphaFold Protein Structure Database, making freely available structural predictions for all proteins in the human proteome (Jumper *et al*, 2021). The AlphaFold aligorithm predicts that the NZF domain of Sharpin contacts an interface between the PH-like domain and the UBL domain (**Fig. 4F**), so assuming the structural prediction is correct, this region may form the amylose-binding interface, and removal of the NZF domain may increase accessibility to amylose to enhance binding.

### Allosteric activation of HOIL-1 by Met1-linked and Lys63-linked ubiquitin oligomers

The experiments described in the preceding section suggested a role for HOIP in recruiting HOIL-1 to glycogen. We therefore wondered whether HOIP might also facilitate the HOIL-1-catalysed ubiquitylation of glycogen in other ways. HOIL-1 contains an Npl4 Zinc Finger (NZF) domain that is reported to bind M1-Ub dimers about 10-fold more strongly than K63-Ub dimers (Sato *et al*, 2011). We therefore wondered whether M1-Ub oligomers, formed by the action of HOIP, might bind to the NZF domain of HOIL-1, inducing a conformational change that accelerated the HOIL-1-catalysed ubiquitylation of maltoheptaose. We found that the inclusion of M1-Ub or K63-Ub dimers (**Fig. 5A)** or tetramers (**Fig. 5B**) did indeed speed up the rate of conversion of maltoheptaose to ubiquitylated-maltoheptaose. Remarkably, after only one minute the conversion of ubiquitin to mono-ubiquitylated maltoheptaose was already greater than after 2 h in the absence of M1-Ub or K63-Ub oligomers. After 10-15 minutes nearly all the ubiquitin in the assays had been converted to ubiquitylated maltoheptaose (**Figs. 5A and 5B**). The experiment demonstrated that the rate of ubiquitylation of maltoheptaose was accelerated at least 100-fold in the presence of small M1-Ub or K63-Ub oligomers.

**Figure 5.**
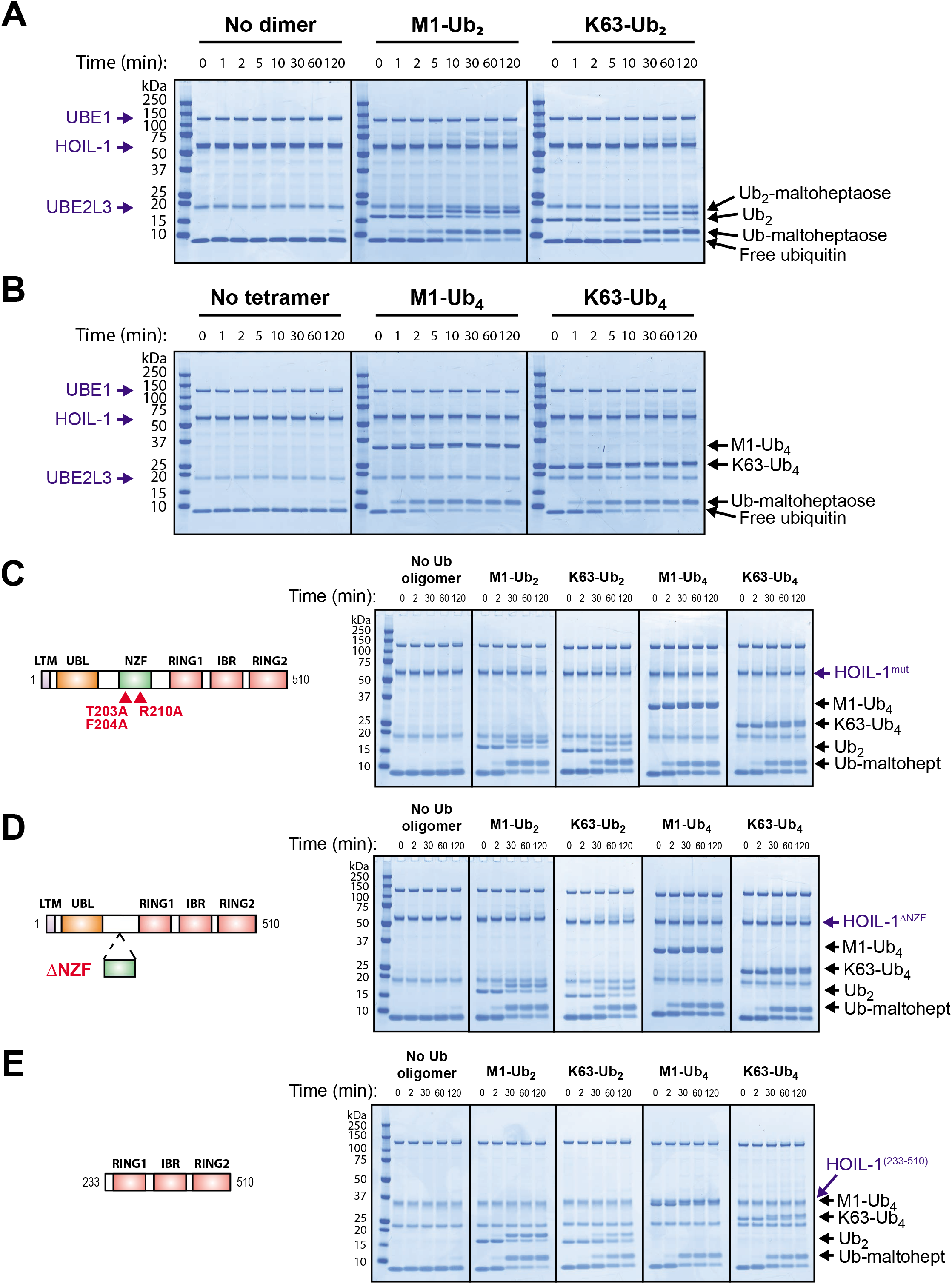
HOIL-1 is allosterically activated by Met1- and Lys63-linked ubiquitin chains. **(A)** Maltoheptaose ubiquitylation by HOIL-1 was assayed at 30°C in the presence of Met1- or Lys63-linked ubiquitin dimers for the indicated times and visualised by Coomassie staining. **(B)** Maltoheptaose ubiquitylation by HOIL-1 was assayed at 30°C in the presence of Met1- or Lys63-linked ubiquitin tetramers for the indicated times and visualised by Coomassie staining. **(C)** HOIL-1[T203A/F204A/R210A] was assayed at 30°C in the absence or presence of the indicated ubiquitin dimers and tetramers. These mutations have been reported to impair the binding of ubiquitin to the NZF domain (Gomez-Diaz *et al*., 2021; Sato *et al*., 2011) and their approximate location in HOIL-1 is indicated (schematic not to scale). **(D)** Assay of HOIL-1[Δ194-222] lacking the NZF core domain at 30°C in the absence or presence of the indicated ubiquitin oligomers. **(E)** Allosteric activation of HOIL-1[233-510] (lacking the LTM, UBL and NZF domains at the N-terminus) by ubiquitin oligomers.

To investigate whether the M1-Ub and K63-Ub oligomers exerted their effects on HOIL-1 activity by binding to the NZF domain, we made mutations in this domain that have been reported to impair the binding of M1-Ub dimers (Gomez-Diaz *et al*, 2021; Sato *et al*., 2011). Surprisingly, these mutations did not reduce the rate of ubiquitylation of maltoheptaose by either M1-Ub or K63-Ub oligomers **(Fig. 5C)**. To check this result, we next made a mutant that removed most of the NZF domain, a deletion mutant that is known to abolish the binding of M1-Ub dimers to the NZF domain (Sato *et al*., 2011). However, the activation of HOIL-1 by M1-Ub and K63-Ub oligomers remained unimpaired (**Fig. 5D**). Moreover, the N-terminally truncated HOIL-1[233-510] mutant, which lacks the LUBAC-tethering motif (LTM), UBL and NZF domains was also activated by M1-Ub and K63-Ub oligomers similarly to wild type HOIL-1 (**Fig. 5E**). Taken together, these experiments revealed that the small Ub-oligomers did not activate HOIL-1 by binding to the NZF domain or any of the other domains located in the N-terminal region of the protein. They also suggested that HOIL-1 contained another binding site(s) for M1-Ub and K63-Ub oligomers that was most likely located within the RBR domain itself.

The related RBR E3 ligase family members HOIP and Parkin are allosterically activated by the binding of M1-Ub dimers (HOIP) and Ser65-phosphorylated ubiquitin (Parkin) to a ubiquitin-binding site within the RBR domain (Lechtenberg *et al*, 2016; Wauer *et al*, 2015). We therefore wondered whether the allosteric binding site for M1-Ub dimers and K63-Ub dimers in HOIL-1 might be located in the analogous region of the protein and made mutations equivalent to those that impair activation of HOIP and Parkin by M1-Ub dimers and phospho-ubiquitin respectively **(Fig. 6A)**. Mutation of Ile358 to Arg (equivalent to the A320R mutation in Parkin that abolishes phospho-Ub binding) or the double mutation F381A/E383A (equivalent to combining the I807A and E809A mutants that disrupt the allosteric activation of HOIP) resulted in a pronounced reduction or loss of HOIL-1 activation by Ub dimers, but had less effect on the activation of HOIL-1 by K63-Ub tetramers and very little effect on ligase activation by M1-Ub tetramers (**Fig. 6B-D**).

**Figure 6.**
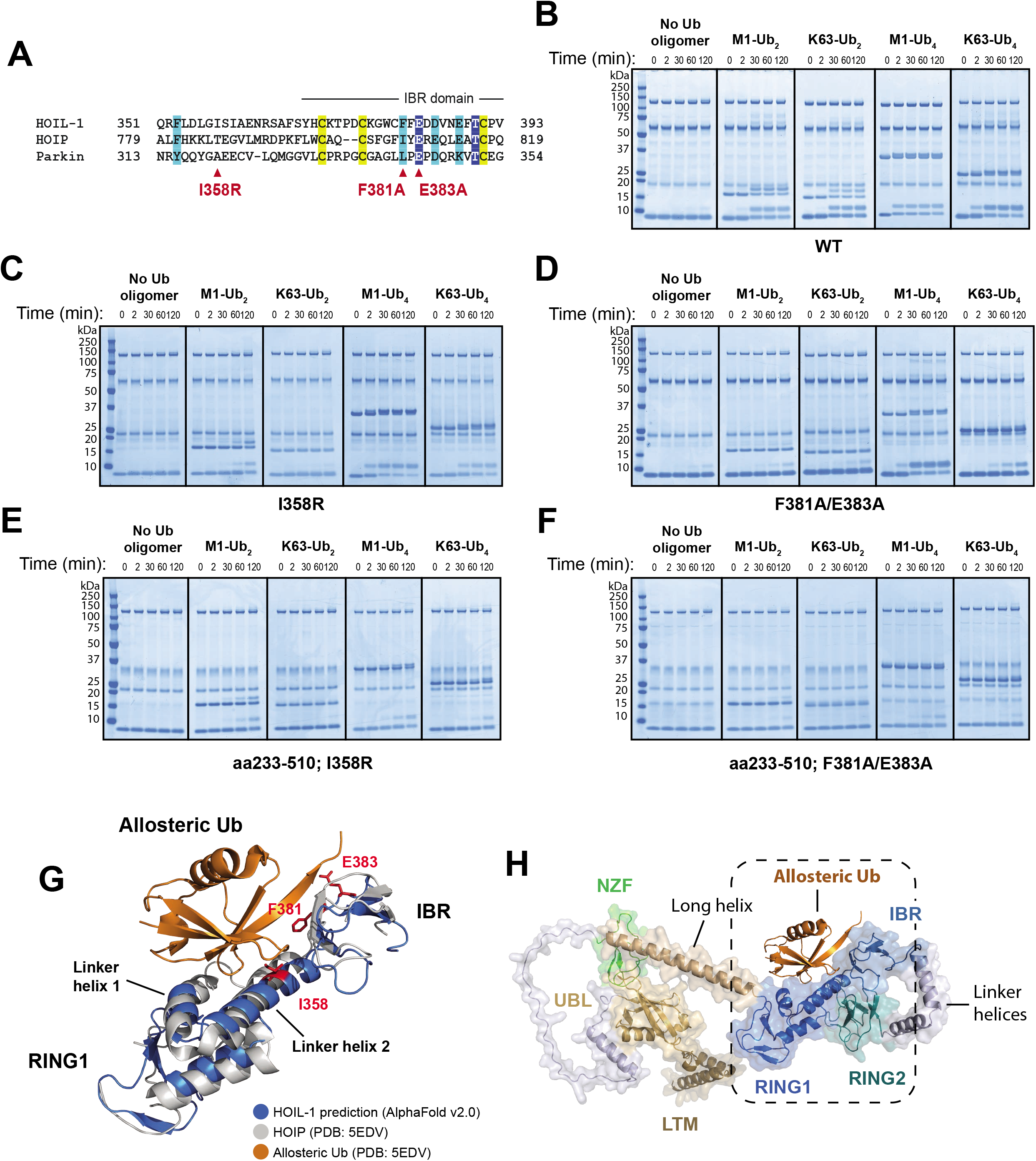
Allosteric ubiquitin binding to the IBR and preceding helical region are important for HOIL-1 activation. **(A)** Amino acid sequence alignment of HOIL-1, HOIP and Parkin centred at the start of the IBR domain. Conserved zinc coordinating residues are highlighted in yellow, other highly conserved residues are highlighted with dark blue, and weaker conservation of property is indicated by light blue highlight. Red triangles denote the position of the mutations indicated. **(B)** Time-course assay performed at 30°C featuring wild-type HOIL-1 in the absence or presence of the indicated ubiquitin oligomers, visualised by Coomassie staining. **(C)** HOIL-1[I358R], assayed as in (B). **(D)** HOIL-1[F381A/E383A], assayed as in (B). **(E)** The truncated HOIL-1[233-510; I358R] mutant was assayed as in (B). **(F)** The truncated HOIL-1[233-510; F381A/E383A] mutant was assayed as in (B). **(G)** The sequence from RING1-IBR was aligned by overlaying the HOIL-1 AlphaFold prediction (residues 282-404) with residues 699-827 in the crystal structure of HOIP-RBR (PDB: 5EDV). The location of the allosteric ubiquitin from that HOIP structure is also shown. HOIL-1 residues I358, F381 and E383 are highlighted in red. **(H)** Full-length HOIL-1 structural prediction with likely location of allosteric ubiquitin shown. The region shown in (G) is indicated.

We next combined these RBR domain mutations with the HOIL-1[233-510] truncation, which ablated the activation of HOIL-1 by ubiquitin tetramers almost completely (**Figs. 6E and 6F**). This suggested that, while not essential for the allosteric activation of HOIL-1, a region(s) N-terminal to the RBR may contribute to the binding of longer Ub oligomers. In order to gain more insight into how ubiquitin oligomers influence HOIL-1 activity, we examined the HOIL-1 structure predicted in the AlphaFold Protein Structure Database. Alignment of AlphaFold’s HOIL-1 prediction with the experimentally-derived structure of HOIP (Lechtenberg *et al*., 2016) revealed remarkable similarities in the conformations of the RING1 and IBR domains of these two proteins (**Fig. 6G**), and suggested that the allosteric ubiquitin is accommodated by a common binding site in HOIL-1 and HOIP (**Figs. 6G** and **6H**). Our mutagenesis data also revealed that hydrophobic interactions with ubiquitin’s Ile44 hydrophobic patch, coupled with the use of a glutamic acid “clamp” to bind the two arginines (R72/R74) at the C-terminus of ubiquitin are employed similarly in both proteins. Using this ubiquitin binding site as a reference point, we modelled the binding of tetra-ubiquitin to HOIL-1. This suggested an attractive model whereby linear ubiquitin tetramers wrap around the RBR domain, potentially making contacts with the RING2 domain that hold it in position close to the E2-binding RING1 domain (**Fig. EV3A**). While this model accommodates ubiquitins two, three and four of the tetramer, it leaves the first ubiquitin rather unaccounted for. To address this, we compared the AlphaFold-derived HOIL-1 protein with structural predictions produced by RoseTTAFold, a different protein-structure prediction software (Baek *et al*, 2021). AlphaFold and RoseTTAFold disagree on the likely angle at which the predicted long central helix in HOIL-1 lies relative to the RING1 domain (**Fig. EV3B**). Intriguingly, RoseTTAFold predicts the position of this central helix to lie adjacent to the Ub-1 that is unbound in the AlphaFold prediction (**Figs. EV3C and EV3D**). Therefore, Ub tetramer-binding potentially bridges and connects all of the component RBR domains, as well as the helical region N-terminal to these.

### HOIL-1 can polyubiquitylate maltoheptaose by catalyzing the *en bloc* transfer of Ub-oligomers directly to the oligosaccharide

Interestingly, during the HOIL-1 catalysed reaction, the M1-Ub or K63-Ub dimers and tetramers included to accelerate the rate of formation of monoubiquitylated maltoheptaose, themselves became attached covalently to maltoheptaose, and at a similar rate to the formation of monoubiquitylated maltoheptaose (**Figs. 5 and 6**). This *en bloc* transfer of pre-assembled ubiquitin chains to maltoheptaose is dependent on UBE1 and UBE2L3, and did not require free mono-ubiquitin **(Figs. 7A and 7B)**. The oxyester bond linking maltoheptaose to the ubiquitin dimer or tetramer was hydroxylamine-sensitive and could not be cleaved by the M1-Ub-specific deubiquitylase Otulin or the K63-Ub-specific deubiquitylase AMSH-LP **(Figs. 7C and 7D)**. Performing time-course assays allowed the visualisation of a dithiothreitol-sensitive tetra-ubiquitin adduct on UBE1, UBE2L3 and HOIL-1 that was prominent at early timepoints but decreased after 2 min **(Fig. 7E left panel)**. When wild-type HOIL-1 was replaced with a catalytically inactive C460S or C460A mutant the transient adducts on UBE1 and UBE2L3 could not be discharged and persisted, while the use of the C460S mutant similarly stabilised a dithiothreitol-insensitive oxyester-linked tetraubiquitin adduct on HOIL-1 **(Fig. 7E, middle and right-hand panels)**. These experiments are consistent with the HOIL-1-catalysed conjugation of ubiquitin oligomers to maltoheptaose *en bloc*. The HOIL-1-initiated polyubiquitylation of glycogen (and proteins) in cells, might therefore be catalysed in a single step, and not sequentially by HOIL-1 catalysed monoubiquitylation followed by the addition of further ubiquitin molecules to monoubiquitylated glycogen through the action of other E3 ligases.

**Figure 7.**
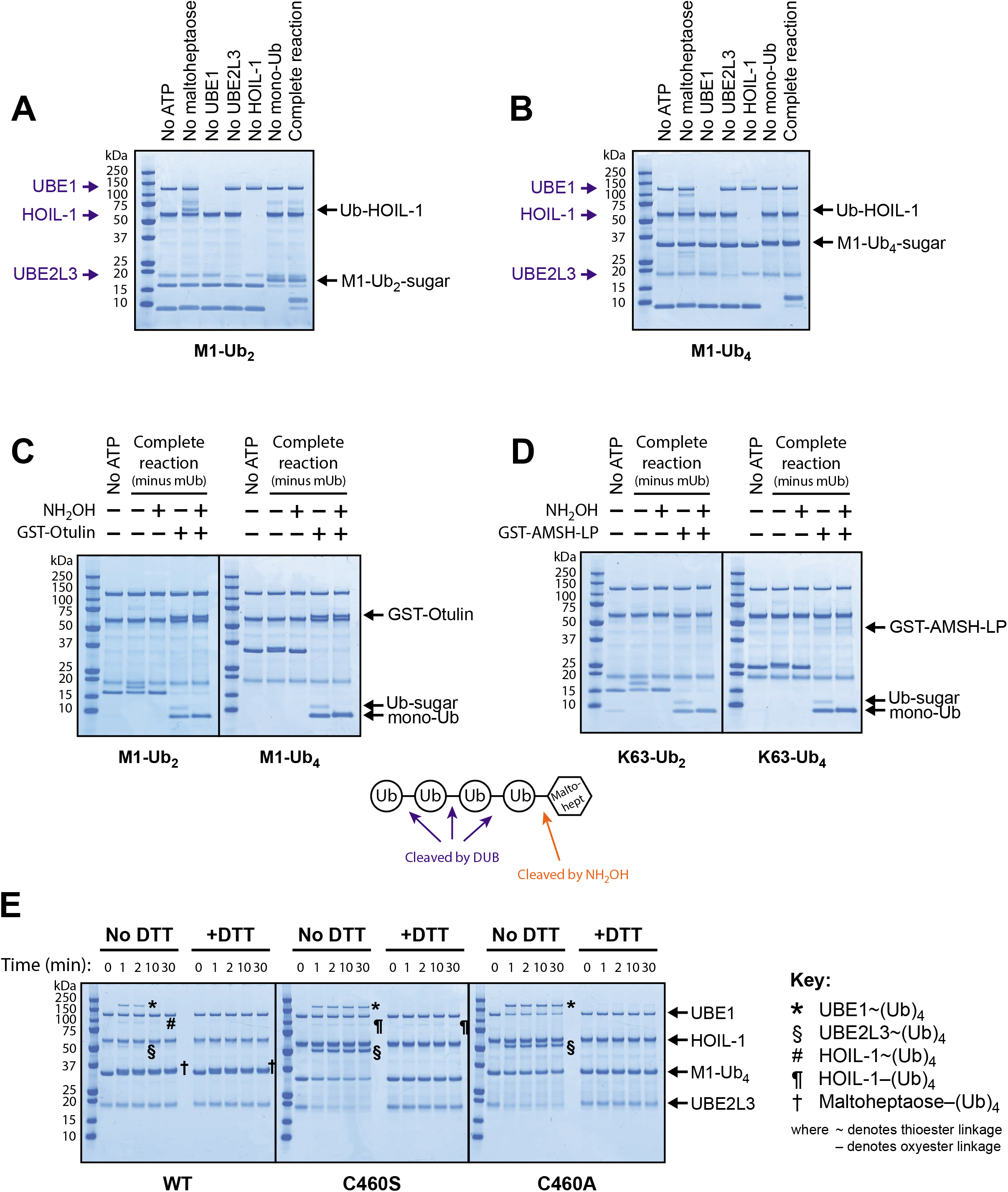
HOIL-1 catalyses *en bloc* transfer of pre-formed ubiquitin oligomers directly to maltoheptaose. **(A)** Requirements for *en bloc* transfer of linear ubiquitin dimer to maltoheptaose were investigated by leaving out the indicated reaction components. **(B)** Requirements for *en bloc* transfer of linear ubiquitin tetramer to maltoheptaose were investigated by leaving out the indicated reaction components. **(C)** *En bloc* transfer of linear ubiquitin dimers and tetramers was performed for 45 min at 37°C in the absence of monomeric ubiquitin. The reaction was terminated (and further reaction prevented) by addition of the UBE1 inhibitor MLN7243 to a final concentration of 25 μM and incubation for 15 min at 30°C. This was followed by treatment with 1.3 M hydroxylamine and/or 1 μM GST-Otulin for 60 min at 37°C. Reaction products were separated by SDS-PAGE and visualised by Coomassie staining. **(D)** *En bloc* transfer of K63-linked ubiquitin dimers and tetramers was performed in the absence of monomeric ubiquitin, followed by inhibition with 25 μM MLN7243 and treatment with 1.3 M hydroxylamine and/or 0.2 μM GST-AMSH-LP[264-436]. **(E)** *En bloc* transfer of M1-Ub_4_ to maltoheptaose was performed in the absence of monomeric ubiquitin using wild-type HOIL-1 or the catalytically inactive mutants C460S and C460A. Reactions were terminated at the indicated timepoints by the addition of NuPAGE LDS gel loading buffer without or with 50 mM dithiothreitol (DTT), subjected to SDS-PAGE, and visualised by Coomassie staining. DTT-sensitive adducts indicate thioesters whereas those insensitive to reducing agent represent oxyesters, such as that in maltoheptaose–(Ub)_4_ or the engineered oxyester linkage between M1-Ub_4_ and Ser460 in HOIL-1[C460S].

## Discussion

Ever since the discovery of lysosomal α1:4 glucosidase deficiency (Pompe’s disease (Hers, 1963)) it has been known that glycogen-like molecules are continually being transported from the cytosol to the lysosomes, where they are taken up and hydrolysed by lysosomal α1:4 glucosidase enabling the glucose units in glycogen to be recycled. More recently, it has been recognised that this process occurs by a mechanism akin to autophagy and has therefore been termed glycophagy (Zhao *et al*, 2018). In this process, it has been proposed that starch-binding domain-containing protein 1 (Stbd1), a protein localised to perinuclear or lysosomal membranes, acts as a cargo receptor to recruit glycogen to lysosomes (Jiang *et al*, 2010).

Autophagy requires the ubiquitylation of organelles or pathogenic bacteria, prior to their uptake into endo-lysosomes where they are destroyed (Noad *et al*, 2017; Polajnar *et al*, 2017; van Wijk *et al*, 2017), and the possibility that glycophagy requires the ubiquitylation of glycogen-associated proteins has been considered by others (Brewer & Gentry, 2019; Sanchez-Martin *et al*, 2020). However, glycophagy has generally been considered to be a device for removing normal glycogen from cells to prevent its accumulation in excess of the cell’s requirements, enabling the excess glucose to be recycled for use by other cells (Zhao *et al*., 2018). However, it is possible that instead, or in addition, glycophagy is a quality control mechanism that functions to remove abnormal forms of glycogen from cells, for example glycogen molecules with few α1:6 branch points that may normally be formed adventitiously in trace amounts by errors of metabolism. If not removed rapidly, these starch-like molecules precipitate as polyglucosan deposits, which gradually accumulate in tissues until they reach levels that cause serious damage to tissue functions as shown, for example, by the fatal diseases that arise in humans and mice with deficiencies in glycogen branching enzyme, lysosomal α1:4 glucosidase or HOIL-1.

Until now, the quality control mechanism by which unbranched glucose polymers can be distinguished from normal glycogen has been obscure but, based on the results presented in this paper, we suggest the following new working hypothesis. We suggest that α1:4-linked oligosaccharides that have failed to form α1:6 branch points when they attain a certain length may then undergo HOIL-1 catalysed ubiquitylation that triggers their recognition by the glycophagy machinery, enabling their uptake into lysosomes and hydrolysis. Ubiquitylation might also enhance the solubility of unbranched glycogen preventing its precipitation until uptake into lysosomes has taken place.

HOIL-1 is associated with HOIP and Sharpin in LUBAC and, in the present study we found that both HOIP and Sharpin interact with amylose *in vitro*, suggesting that one of their functions may be to target HOIL-1 to glycogen in cells. We also found that small M1-Ub oligomers produced by HOIP are powerful allosteric activators of the HOIL-1-catalysed ubiquitylation of maltoheptaose, indicating a second way in which HOIP may facilitate the ubiquitylation of unbranched α1:4-linked glucosaccharides. Interestingly, these studies revealed that the allosteric binding site for these M1-Ub or K63-Ub dimers and tetramers was not the previously described M1-Ub-binding NZF domain of HOIL-1, but a region within the RBR domain.

In conclusion, our results are the first documented example of sugar ubiquitylation, introducing a new aspect to ubiquitylation research. Recent work from the Randow laboratory has described the ester-linked ubiquitylation of bacterial lipopolysaccharide by the E3 ligase RNF213 (Otten *et al*, 2021). However, the authors did not identify the site of ubiquitin attachment, so it is unclear whether it is the lipid or sugar moieties of lipopolysaccharide that undergo ubiquitylation. Nevertheless, their paper and the present study, indicate that ester-linked ubiquitylation of non-proteinaceous biological molecules is a tactic employed by more than one family of ubiquitin ligases to enable the ubiquitylation of amine-free substrates

## Experimental Procedures

### Histological examination of mouse tissues

E3 ligase-inactive HOIL-1[C458S] mice (Kelsall *et al*., 2019) and wild type littermates were euthanized by injecting a lethal dose of sodium pentobarbitone. The brain, heart, lungs, and liver were removed and fixed for 48–72 h in 10% neutral buffered formalin. Tissues were processed and 3 μm deparaffinized sections were treated for 45 min with 0.5% (w/v) α-amylase (Sigma), rinsed for 30 min in water and processed for Periodic Acid Schiff (PAS) staining. The tissue sections were assessed by a veterinary pathologist (C.S.) blinded to the genotype of the mice in the different cohorts. A simple semiquantitative scoring system was used for the assessment of α-amylase resistant-PAS positive materials. Scores for amylase-resistant PAS-positive deposits were defined as:- (1) occasional granular deposits, just detectable by light microscopy: (2) increased numbers of deposits, readily detectable at x10 magnification; (3) markedly increased numbers of granular deposits easily identifiable by light microscopy. The photomicrographs were captured with Nanozoomer software from whole slide image scans prepared using a Hamamatsu slide scanner.

### Plasmids and proteins

DNA constructs were generated by MRC-PPU Reagents & Services and sequenced by MRC-PPU DNA Sequencing & Services (www.dnaseq.co.uk). All plasmids used in this study are available upon request at https://mrcppureagents.dundee.ac.uk. Full length untagged HOIL-1 (and various mutants thereof) was expressed as His_6_-SUMO-HOIL-1 in BL21(DE3) *E. coli* cells before removal of the tag with the SUMO protease Ulp1 and buffer exchange by size exclusion chromatography or dialysis into PBS, 1 mM TCEP. Human ubiquitin, GST-Otulin and GST-AMSH-LP[246-436] were expressed and purified as described elsewhere (Ritorto *et al*, 2014). The expression and purification of His_6_-UBE1 and UBE2L3 (Kelsall *et al*., 2019), GST-CHIP (Zhang *et al*, 2005), and HHARI (Duda *et al*, 2013) have been described previously. HOIP and Sharpin were expressed in BL21 cells as GST-tagged fusion proteins, purified on GSH-Sepharose and collected by elution with 10 mM glutathione or by removal of the GST-tag on the resin using PreScission Protease. MBP-His_6_ was purified by Ni^2+^-Sepharose chromatography. Lys63-linked and linear di- and tetra-ubiquitin chains were produced and purified according to previously reported methods (Dong *et al*, 2011; Komander *et al*, 2008). All proteins were prepared in PBS, 1 mM TCEP and stored in aliquots at −80°C.

### HOIL-1 catalysed ubiquitylation assays using fluorescent Cy5-labelled ubiquitin

Reactions (20 μL) contained 500 nM His_6_-UBE1, 2 μM UBE2L3, 2 μM HOIL-1, 10 μM Cy5-ubiquitin (South Bay Bio) and 10 mg/mL bovine liver glycogen (Sigma) in Phosphate Buffered Saline pH 7.4 containing 0.5 mM TCEP and 5 mM Mg^2+^-ATP. Reactions were incubated at 37°C for 1 h with gentle mixing. Where indicated, reactions were then incubated in the presence of 50 μg/mL (~0.1 units) human salivary α-amylase (Sigma) or 1.5 M hydroxylamine for a further 60 min at 37°C. Reactions were stopped by adding NuPAGE LDS sample loading buffer supplemented with 50 mM DTT, and denatured at room temperature for 10 min. Reaction products were separated by gel electrophoresis on 4-12% Bis-Tris gradient gels using MES SDS running buffer (Invitrogen) and stained with Flamingo fluorescent protein stain (Bio-Rad). Visualisation of the fluorescent signals was performed using the ChemiDoc MP Imaging System (Bio-Rad) and ImageJ software. In the case of assays using glycogen as substrate, these gels were then stained with Periodic acid-Schiff stain using the Pierce Glycoprotein Staining Kit (Thermo Scientific) and Coomassie stained with InstantBlue Protein Stain (Expedeon).

### HOIL-1 catalysed ubiquitylation assays (non-fluorescent)

Reactions (typically 10-20 μL) contained 500 nM His_6_-UBE1, 2 μM UBE2L3, 2 μM HOIL-1, 10 μM ubiquitin and 20 mM maltosaccharide substrate (unless otherwise indicated) in Phosphate Buffered Saline pH 7.4, 0.5 mM TCEP and 5 mM Mg^2+^-ATP. Reactions were incubated at 37°C for 1 h with gentle mixing, unless otherwise stated. Where indicated, reactions were then incubated in the presence of 50 μg/mL (~0.1 units) human salivary α-amylase or 1.5 M hydroxylamine for a further 60 min at 37°C. Reactions were quenched by addition of NuPAGE LDS sample loading buffer supplemented with 50 mM DTT, and denatured at room temperature for 10 min. Reaction products were separated by gel electrophoresis on 4-12% Bis-Tris gradient gels using MES SDS running buffer and coomassie stained with InstantBlue Protein Stain before analysis on the ChemiDoc MP Imaging System. Maltoheptaose was purchased from CarboSynth. Maltose was from Sigma. Cyclomaltoheptaose (β-cyclodextrin) and 6-mercapto-cyclomaltoheptaose (heptakis-(6-deoxy-6-mercapto)-β-cyclodextrin) were from AraChem. The UBE1 inhibitor MLN7243 was purchased from Active Biochem.

Where noted, HOIL-1 activity was stimulated by addition of chain-type specific ubiquitin dimers or tetramers at a concentration of 2 μM. In these experiments the ubiquitin chains were pre-incubated for 30 min at 30°C with all other reaction components except ATP. Reactions were initiated by adding Mg^2+^-ATP to a final concentration of 5 mM. In order to slow the reaction rate and allow better visualisation of the differences between ligase activation in the presence of different chain types, maltoheptaose concentration was dropped to 10 mM and reactions proceeded at the lower temperature of 30°C for the indicated times.

### Preparation of purified Ub-maltoheptaose

Ubiquitylation reactions were carried out as described previously and terminated through the addition of 20% (v/v) trifluoroacetic acid (TFA) to a final concentration of 2%. Ubiquitylated-maltoheptaose was separated from unconjugated free ubiquitin by reverse phase-high performance liquid chromatography (RP-HPLC) using a Dionex Ultimate 3000 System. A Thermo BioBasic column (250 x 10 mm) was equilibrated in aqueous buffer containing 20% (v/v) acetonitrile supplemented with 0.1% (v/v) TFA. A flow rate of 2.3 mL/min and an increasing gradient of acetonitrile from 20% to 60% over 60 min was utilised for sufficient separation. Separated fractions were validated using MALDI-TOF mass spectrometry before being pooled and freeze-dried.

### MALDI-TOF mass spectrometry

For MALDI-TOF sample preparation a mixture containing a 1:1 ratio of 2% TFA (v/v) and 2,5 Dihydroxyacetophenone (DHAP) matrix solution (7.6mg of 2,5 DHAP in 375 μL 100% ethanol and 12 μL of 12 mg/mL diammonium hydrogen citrate) was added to the sample in a 1:1 ratio. 1 μL of the solutions were spotted in duplicate onto an MTP AnchorChip 1,536 Target (Bruker Daltronics). Samples were air dried at room temperature prior to analysis. All spectra were acquired using a Rapiflex MALDI-TOF mass spectrometer (Bruker Daltronics) equipped with Compass 1.3 control. Recording took place in automatic mode (AutoXecute, Bruker Daltronics), allowing 6-9 seconds per target spot as described previously (De Cesare *et al*, 2020). Spectra were visualised using FlexControl software and processed by FlexAnalysis software (version 4.0).

### NMR spectroscopy

[*U*-^15^N,^13^C]-labelled Ubiquitin was produced by growing transformed *E. coli* BL21 (DE3) cells (New England Biolabs) in M9 enriched medium supplemented with ^15^NH_4_Cl (1g/L) and ^13^C-glucose (2 g/L). All spectra were recorded on a Bruker Avance III HD 800 MHz equipped with triple-resonance cryoprobe. Purified [*U*-^15^N,^13^C]-labelled ubiquitin-maltoheptaose conjugate was freeze dried and dissolved in 0.5 mL of 20 mM phosphate buffer pH 6.5, 150 mM NaCl, 10% D_2_O to a final concentration of 0.5 mM. The sequential NMR assignment of ubiquitin was amended using HN(CO)CACB, HNCACB, HN(CA)CO and HNCO triple resonance spectra were acquired at 303 K (Cavanagh *et al*, 2007). 2D heteronuclear spectra such as ^1^H-^13^C Heteronuclear Multiple Bond Correlations (HMBC) spectroscopy and Heteronuclear single quantum correlation (HSQC) were recorded with standard experiments.

### Glucan binding assay

Amylose covalently bound to agarose resin (New England Biolabs) was pre-incubated with 10 mg/mL bovine serum albumin (BSA) for 30 min with mixing a room temperature to block non-specific binding. The resin was pelleted by centrifugation (500 × g for 2 min) and washed three times with 10 mM Phosphate Buffer, pH 7.4, containing 137 mM NaCl, 1 mM TCEP, 0.01% (v/v) Tween-20 and 0.01 mg/mL BSA (Binding Buffer). 1 μg of bacterially-expressed protein was then incubated with 30 μL of this blocked amylose resin with mixing for 60 min at 4°C in a final volume of 150 μL of binding buffer. Following incubation, the beads were pelleted by centrifugation (500 × g for 2 min), the supernatant removed and retained, and the resin washed three times with binding buffer before finally resuspending with buffer to the original 150 μL starting volume. Bound proteins were eluted by addition of NuPAGE LDS gel loading buffer and proteins in the supernatant and pellet were analysed by gel electrophoresis followed by western blotting using the following antibodies: anti-HOIP (#SAB2102031) and anti-HOIL-1 (#HPA024185) were from Sigma-Aldrich; anti-MBP (#E8032S) was from New England Biolabs; HRP-conjugated anti-rabbit secondary (#7074), and HRP-conjugated anti-mouse secondary (#7076) were from Cell Signaling Technology; and HRP-conjugated anti-sheep secondary (ab97130) was from Abcam. Polyclonal antibodies against GST (#S902A), CHIP (#S471B) and HHARI (#S622D) were generated by MRC-PPU Reagents & Services and are available upon request at https://mrcppureagents.dundee.ac.uk.

## Acknowledgements

This study was supported by Wellcome Trust Investigator Award 209380/Z/17/Z (to PC). We thank our colleagues Ron Hay, Michael Ferguson, Yogesh Kulathu, Sven Lange, Mathieu Soetens and Satpal Virdee for helpful suggestions and advice during the course of this project.

## Author contributions

IRK and PC conceived the study. The experiments were performed by CLS and SKN (Figs 1 and EV1), PMG, EHM and IRK (Fig 2), EHM (Fig EV2), YX, SJM and EHM (Fig 3) and IRK (Figures 4–7 and EV3). AK, IRK and EHM produced bacterially expressed proteins and NTW produced all the DNA constructs. PC, IRK, EHM and SJM wrote the paper.

## Expanded View Figures

**Figure EV1.**
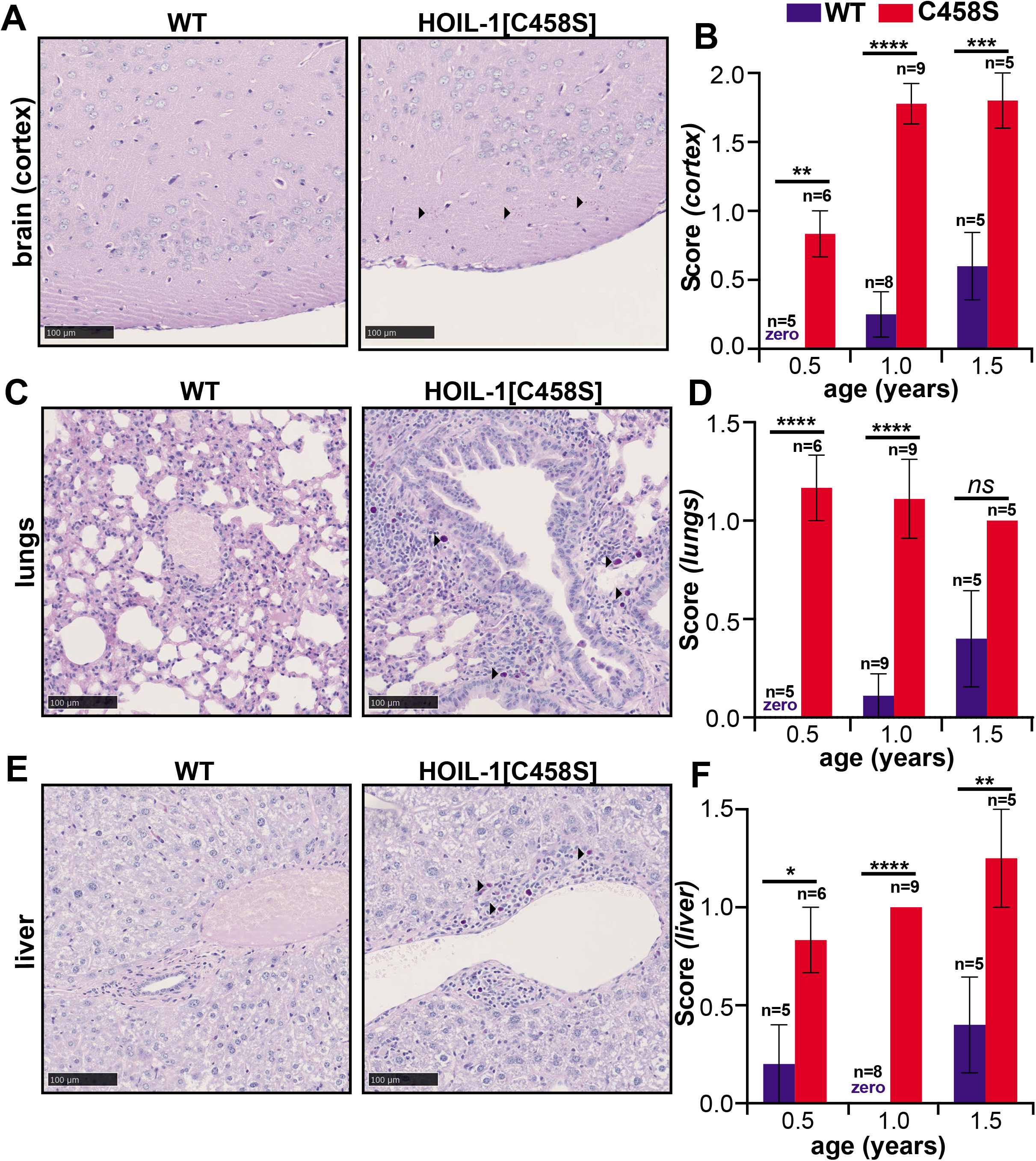
Deposition of α-amylase-resistant polyglucosan deposits in the brain cortex, lungs and liver of HOIL-1[C458S] mice. **(A, C, E)** Representative PAS-stained sections of the brain cortex (A) lung (C) and liver (E) of one year old HOIL-1[C458S] and WT mice are shown. Scale bar = 100 μm. Arrow heads indicate α-amylase-resistant PAS-positive polyglucosan deposits. **(B, D, F)** Graphs quantitating α-amylase resistant PAS scores of the brain cortex (B) lung (D) and liver (F) of HOIL-1[C458S] (red) and WT (blue) mice aged 0.5, 1.0 and 1.5 years. The number of biological replicates analysed at each age is indicated. The word zero highlighted in blue indicates that no α-amylase resistant, PAS-positive material could be detected in the WT mice. The error bars show mean ± SEM. Statistical significance between the genotypes was calculated by using two-way ANOVA and Šidák’s multiple comparison’s test. *denotes p<0.05, **p<0.01, ***p<0.001 and ****p<0.0001.

**Figure EV2.**
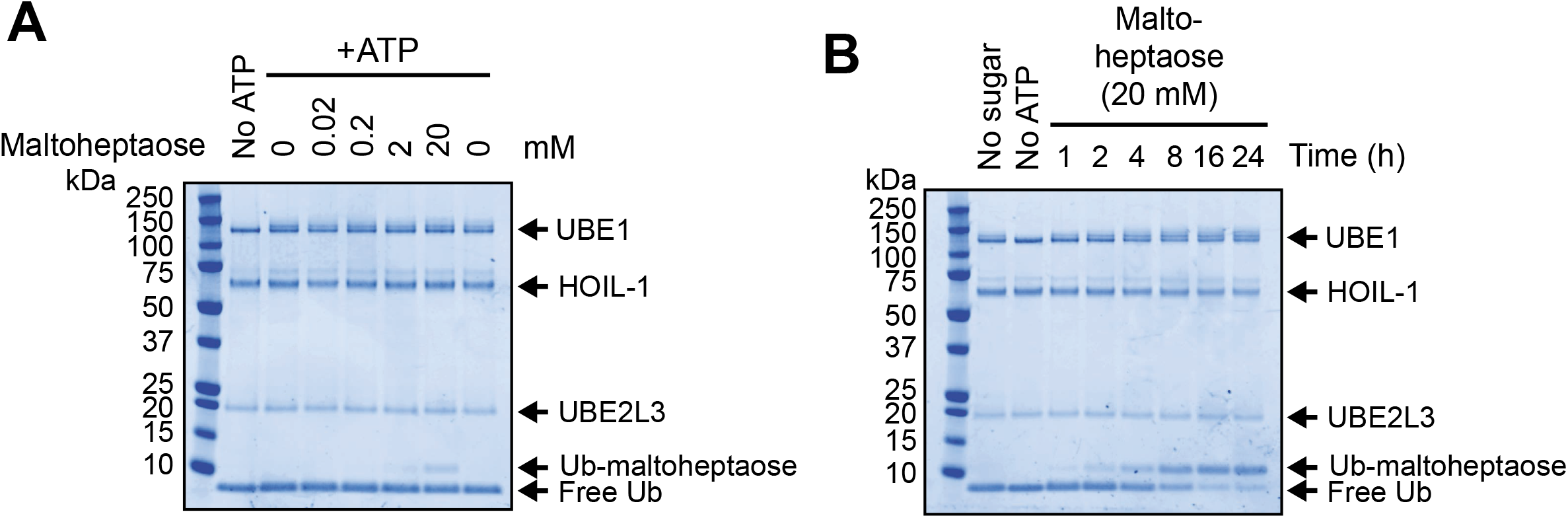
Ubiquitylation of maltoheptaose by HOIL-1. **(A)** Increasing concentrations of maltoheptaose were incubated with bacterially-expressed HOIL-1 for 1 hour at 37°C and reaction products were resolved by reducing SDS-PAGE and visualised by Coomassie staining. **(B)** Maltoheptaose (20 mM) ubiquitylation by HOIL-1 was assayed for the indicated times and visualised by Coomassie staining.

**Figure EV3.**
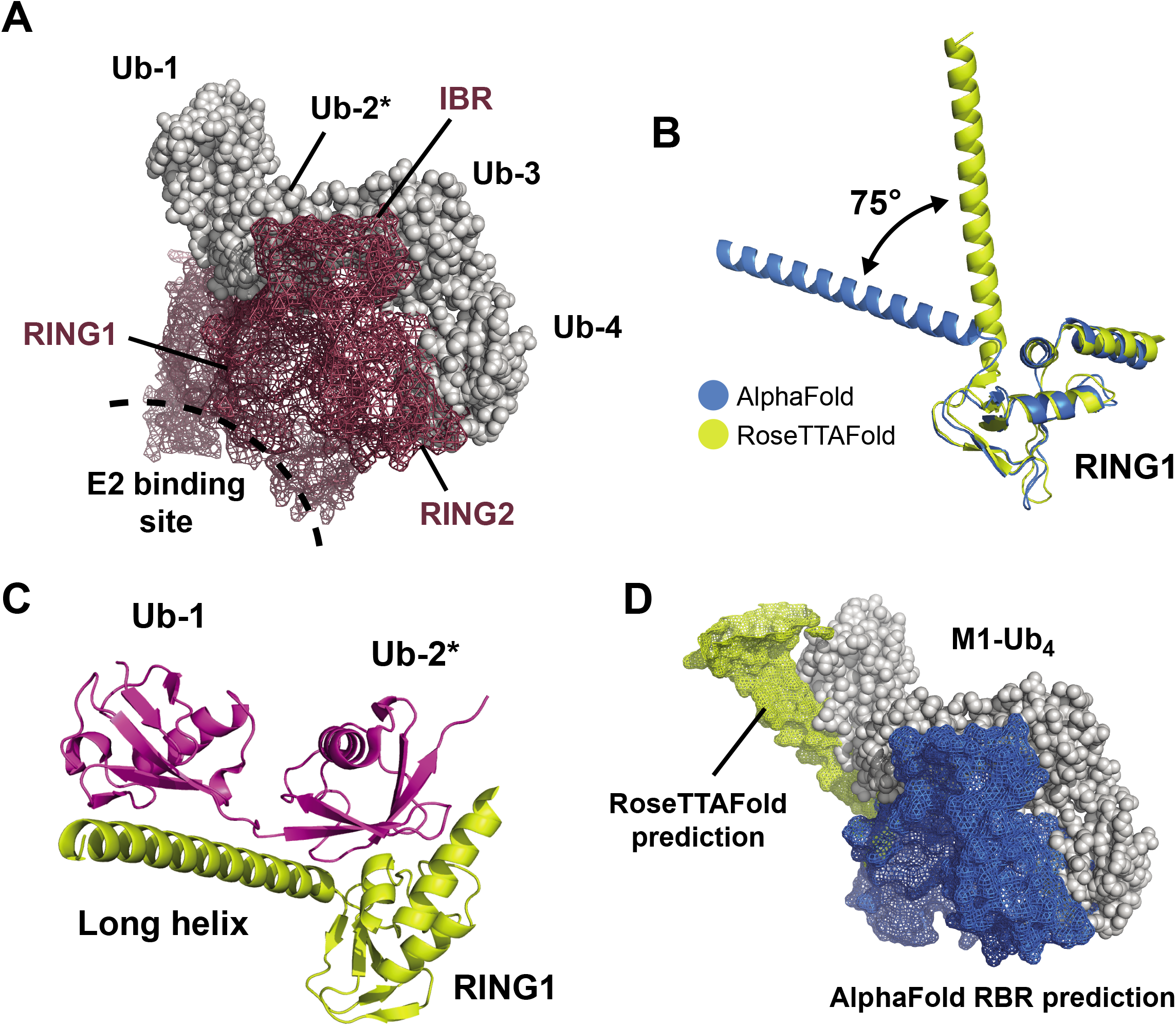
Modelling tetra-ubiquitin binding to HOIL-1. **(A)** Using the model depicted in Figure 6H as a starting point, the structure of M1-Ub_4_ (PDB: 5B83) was modelled onto this to determine where the other ubiquitins in the tetramer might lie. Assuming the already modelled allosteric ubiquitin to represent the second ubiquitin in the tetramer (designated Ub-2* to indicate this) produced the model shown, where the tetramer wraps itself around the RBR domain, making contacts with the IBR and RING2 domains and the helices that sit C-term to the RING1 domain. Ubiquitin is shown as grey spheres and HOIL-1’s surface is represented as brick-red mesh. The approximate location of the E2 binding site is indicated. This structure is rotated approximately 90° on the y-axis relative to Fig 6H. Note that there is a steric clash between ubiquitin-4 and the predicted location of RING2. This suggests that tetramer binding may alter the location of this domain, explaining why Ub_4_-binding influences catalytic activity. **(B)** Comparison of AlphaFold (blue) and RoseTTAFold (yellow) predictions for HOIL-1 residues 233-362. The programs disagree on the angle at which the long helix (residues 233-271) lies relative to the RING1 domain, with a difference of 75° between the two AI-derived predictions. **(C)** The predicted position of helical region 233-271 in the RoseTTAFold-generated structure places it in a much better position to interact with Ub-1 in the ubiquitin tetramer (shown here in magenta). **(D)** AlphaFold’s RBR domain prediction is modelled in blue binding to linear tetra-ubiquitin (grey spheres). Orientation is the same as shown in (A). Helix 233-271 is coloured yellow and presented in the orientation predicted by RoseTTAFold.

